# Local Adaptation and Transcriptomic Plasticity of the Copepod *Acartia tonsa* Under Low Salinity Stress

**DOI:** 10.1101/2025.06.12.659303

**Authors:** Alexandra Hahn, Jennifer C. Nascimento-Schulze, Georgia Avgerinou, Till Bayer, Reid S. Brennan

**Affiliations:** GEOMAR Helmholtz Centre for Ocean Research Kiel

**Keywords:** Local adaptation, osmoregulation, gene expression, copepod, salinity tolerance, plasticity

## Abstract

Salinity—an essential factor shaping marine species distributions—is rapidly shifting due to global change, yet the mechanisms of salinity tolerance and adaptation remain poorly understood. We investigated local adaptation in the calanoid copepod *Acartia tonsa*, a broadly distributed marine species that thrives in the brackish Baltic Sea. Using a common-garden design, we compared physiological and transcriptomic responses to low salinity between populations from the North Sea (>25 PSU) and the Baltic Sea (<15 PSU). Baltic copepods exhibited significantly higher survival under low salinity, indicating local adaptation. While both populations shared a core osmoregulatory strategy involving active ion transport and regulation of amino acids, transcriptomic profiles revealed population-specific differences. Baltic individuals showed a reduced overall gene expression response, yet maintained higher relative expression of osmoregulatory genes—suggesting higher plasticity and a primed response. In contrast, North Sea copepods exhibited broader transcriptional shifts, including downregulation of metabolic and developmental pathways after prolonged stress exposure, possibly reflecting energy conservation mechanisms. These findings reveal that *A. tonsa* possesses both a plastic osmoregulatory strategy and population-level adaptation that enable survival in extreme salinity conditions. While both populations tolerate short-term exposure to low salinity, local adaptation has enhanced the Baltic population’s resilience. This suggests that *A. tonsa* is broadly tolerant of moderate climate-driven salinity declines across most of its distribution. However, our data also indicate potential range contractions in the lowest salinity zones of the Baltic Sea, underscoring the importance of identifying physiological and genetic thresholds in climate resilience studies.

## Introduction

Salinity plays an important role in shaping marine ecosystems as it poses a barrier for dispersal, restricting species distribution. Transitioning from marine conditions to brackish or even fresh water forces marine organisms to either actively transport ions across their cell membrane to maintain a higher internal osmolarity, making them osmoregulators or to lower their osmolarity to be in equilibrium with the environment, making them osmoconformers (Mantel & Farmer, 1983). If an organism is unable to react to decreases in salinity, the resulting influx of water can lead to physiological impairment and potentially death (Lange, 1970). Given this, in habitats with strong salinity gradients such as estuaries, organisms face challenging conditions (Lawrence et al., 2004). When environmental clines, including salinity, are stable over longer time scales, selection pressure can lead populations to adapt to local salinity conditions, giving them a heritable fitness advantage in this specific habitat (Kawecki & Ebert, 2004). Alternatively, when the environment fluctuates within an organisms’ lifetime, plasticity is expected to evolve (Bitter et al., 2021).

Understanding if and how organisms can tolerate environmental stressors such as low salinity, along with assessing their capacity for local adaptation, helps us predict species dispersal, evaluate their potential to invade new environments, and anticipate how organisms may respond to environmental shifts driven by climate change. Though climate change is unanimously regarded as a threat to biodiversity by the scientific community, not all effects are researched equally. While there is extensive literature on ocean warming, ocean acidification, and ocean deoxygenation (Garcia-Soto et al., 2021; Gruber et al., 2021) and the resulting implications for marine life (Deutsch et al., 2024; Leung et al., 2022; Venegas et al., 2023), the consequences of changes in salinity are widely under-researched (Lee, Downey, et al., 2022). Even seemingly minor salinity reductions can push organisms to their tolerance limit and constrain species distribution if environmental salinity drops below the critical salinity threshold separating marine and freshwater organisms (Khlebovich & Abramova, 2000). In both, low and high latitudes, salinity is predicted to decline due to increased river runoff, melting of glaciers and increased precipitation, and changes in ocean fluxes (Du et al., 2019; McCrystall et al., 2021; Skliris et al., 2014). Therefore, understanding the effects of salinity on organisms and populations is crucial for predicting how global change will shape marine habitats.

The copepod *Acartia tonsa* Dana, 1849, is an ideal species to understand local adaptation and responses to changes in salinity. This species is a predominantly coastal calanoid copepod with a global distribution, attributed to its euryhalinity and high thermal tolerance (Cervetto et al., 1999; Svetlichny & Hubareva, 2014; Walter & Boxshall, 2024). This species, and copepods generally, is an integral component of marine food webs, serving as a prey item for commercially important fish species (Mauchline, 1998), linking primary production to higher trophic levels (Turner, 2004). Because *A. tonsa* inhabits coastal environments, it experiences rapid salinity fluctuations within its lifespan and may harbor broad plasticity to tolerate these changes. However, *A. tonsa* also exists along steep salinity gradients, such as the Baltic Sea, a unique ecosystem with a strong east-west salinity gradient from fully marine to nearly fresh water (Szymczycha et al., 2019). Given *A. tonsa*’s rapid generation time, populations likely experience differences in the mean salinity along this cline and may be locally adapted.

In addition to the strong salinity gradient in the Baltic Sea, its enclosed nature also makes it especially vulnerable to climate change impacts (Reusch et al., 2018). While there is some uncertainty in salinity forecasting (Meier et al., 2022), sea surface salinities in the Baltic Sea previously decreased and are predicted to further decrease over the next century (Kankaanpää et al., 2023; Saraiva et al., 2019), posing a challenge for many Baltic species. For a small number of species in the Baltic Sea, including phytoplankton (Pinseel et al., 2023), mussels (Knöbel et al., 2021), and fish (Guo et al., 2016), adaptation to local salinity has been shown, suggesting that some species possess the capacity to adapt to salinity reductions in this region. However, the resilience and adaptive capacity to changing salinity for most species in the Baltic Sea including copepods is still unknown.

Previous transcriptomic analyses aiming to uncover the osmoregulatory strategy of copepods have primarily focused on two species, *Eurytemora affinis* – a calanoid copepod known for its capacity to invade freshwater habitats (Gerber et al., 2016) – and *Tigriopus californicus* – a harpacticoid copepod that inhabits tidal pools subjected to extreme temperature and salinity fluctuations (DeBiasse et al., 2018). However, the mechanisms of osmoregulatory plasticity and presence of salinity local adaptation in widely distributed pelagic copepods remain largely unexplored.

Here, we fill this gap using *A. tonsa* from the Baltic and North Sea to reveal mechanisms of local adaptation and the core of osmoregulatory plasticity. We combined transcriptomics with phenotypic data to understand 1) the conserved osmoregulatory mechanisms of *A. tonsa* enabling tolerance of low salinity stress, 2) whether there is local adaptation to low salinity between populations originating from the brackish Baltic Sea compared to the fully marine North Sea, and 3) the implications for the capacity of *A. tonsa* to tolerate or further adapt to future changes in ocean salinity. By combining transcriptomic data with fitness measurements, we provide a holistic picture of the effects of low salinity on *A. tonsa* from unique osmotic environments.

## Methods

### Sampling and culturing

*Acartia tonsa* populations for this study were collected by 100 μm WP2 net hauls in the North Sea at Wilhelmshaven (WH: 53° 30’46.8’’ N 8° 08’56.4’’E, collected on Sept 09^th^ 2022) hereafter referred to as North Sea, and in the Western Baltic Sea (SW08: 54° 24’46.8’’ N 11° 37’01.2’’ E, collected on Aug 31^st^ 2022) hereafter referred to as Baltic Sea (Fig. 1A). Daily measurements from E.U. Copernicus Marine Service Information (CMEMS) from 1993-2021 showed that salinity conditions were significantly different between these stations (p < 0.001) with the mean surface salinity during the summer months being 26.2 ± 0.9 practical salinity units (PSU) at the North Sea location and 11.3 ± 2.3 PSU at the Baltic sampling station (Fig. S1, data from CMEMS https://doi.org/10.48670/moi-00021). *Acartia tonsa* is only in the water column during the warm summer months (∼ June–October) and we therefore only included measurements from this timeframe. Live animals were sorted to species level and lab cultures started with approximately 200 adult *A. tonsa* individuals. Over days, cultures were gently adjusted to common-garden conditions at 15.5 °C (± 0.3°C) and 15 PSU at 12:12 light:dark regime. The microalgae *Rhodomonas* sp. was kept in the exponential growth phase at the same temperature and salinity as the cultures and fed at concentrations around 500 μgC/L every other day. Common-garden conditions were maintained for at least three generations prior to the experiments. Full water changes were performed approximately every three to four weeks using filtered sea water (using a 0.2 μm filter) and salinity was monitored and adjusted if necessary, using a mix of tap water and deionized water. All experiments were carried out at the culturing temperature of 15.5°C.

**Figure 1.**
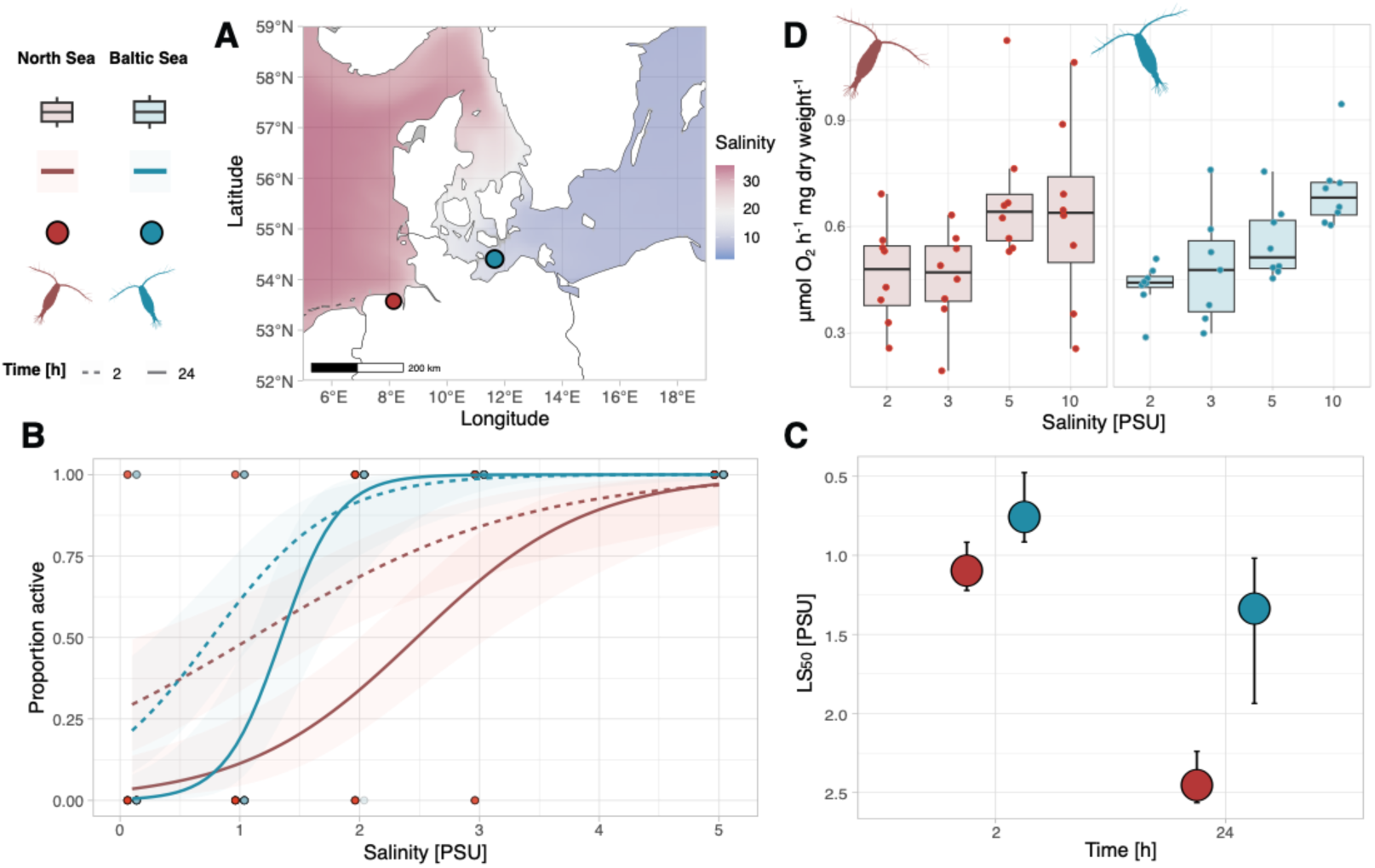
Fitness measurements for Baltic (labeled in blue) and North Sea (labeled in red) *Acartia tonsa* populations. A) Map of sampling locations. Salinity data from CMEMS https://doi.org/10.48670/moi-00021. B) Acute survival of adult females at low salinities for 2 and 24 hours. C) LS50 values for acute survival shown in C, error bars show distribution of replicates D) Oxygen consumption per animal in μmol O_2_ per hour normalized for dried body weight.

### Physiological assays

We conducted a series of experiments to assess physiology and performance of both populations at low salinity conditions: acute mortality of adults, egg production and hatching, respiration, and naupliar survival to adulthood. The data and statistical analysis for all experiments was done in R version 4.2.2 (R Core Team, 2022).

#### Acute mortality

For the acute mortality experiment, 6 salinities were tested (0.1, 1, 2, 3, 5, and 15 PSU). To create the experimental salinities, filtered sea water was diluted with a mix of tap water and deionized water. In total, 90 recently matured females were picked and five at a time were placed in one vial filled with 9 mL of water, so each salinity treatment included three independent replicates with a total of 15 individuals. During the duration of the experiment no food was supplied and survival was monitored after 2 and 24 hours by gently pipetting water up and down. Animals were scored as dead if they did not show any signs of movement to this stimulus. No individuals died at the 15 PSU control salinity. For the analysis, we fit a logistic regression and calculated the salinity at which 50% mortality (LS_50_) was observed for both populations using a generalized linear model with binomial distribution and the *dose.p* function from the *MASS* package (Venables & Ripley, 2002).

#### Egg production and hatching

To measure egg production, recently matured males and females were placed in pairs into 6-well plates filled with 8 mL of filtered sea water at three salinities (5, 10, and 15 PSU) to mate and produce eggs. Each well contained one pair with a total of six replicates per salinity. After 48 hours, adults were removed. After another 48 hours of hatching, the samples were fixed with a drop of 1% Lugol’s solution. The contents were then moved to a petri dish with a counting grid and all intact eggs and nauplii were counted. The number of total eggs laid consisted of all unhatched eggs plus the number of nauplii. Hatching success was defined as the number of nauplii divided by the number of total eggs produced.

#### Respiration rates

To determine base respiration rates, recently matured females were picked and acclimated to the experimental salinities for 16 hours without food. Respiration was measured at two conditions, control (15 PSU) and low salinity (5 PSU). On the day of the experiment, animals were gently rinsed in filtered sea water and then placed in pairs into 80 μL wells filled with filtered sea water at the experimental conditions. To ensure measurements were comparable, both populations were run at the same time and wells assigned at random. Oxygen concentrations were measured for up to 2 hours with a PreSens optical sensor using the SDR_v4.0.0 software. Animals were preserved in 1% Lugol’s solution and later photographed using a Nikon imaging microscope with NIS-Elements software (v. 5.20.000). All pictures were taken with the same magnification to ensure consistency. Prosome length was then measured using the software ImageJ (Schneider et al., 2012). Respiration rates were calculated using the *respR* package (Harianto et al., 2019) and standardized per dry body weight using the length obtained from the images and the conversion factors as defined by Kiørboe et al. (1985). Since Lugol fixation is reported to reduce body size in copepods, we used the factors estimated by Jaspers and Carstensen (2009) to correct for shrinkage, though these effects should be consistent across all individuals.

To assess how acute exposure to extremely low salinity affects respiration, we conducted an additional experiment with even lower salt concentrations and limited acclimation time. Here, we measured the respiration rates of adult females at 2, 3, 5 and 10 PSU. As previous experiments showed that survival is not guaranteed with long term exposure (>24 h) at these extreme low salinities below 5 PSU, we opted for a 2-hour acclimation period at the target salinity for this experiment. Afterwards the procedure was identical to that described above in the first experiment.

#### Survival to adulthood

To test for the effect of low salinity on survival to adulthood, recently matured adults were placed in 100 μm mesh cups inside beakers filled with 10 PSU water. They were allowed to reproduce for 24 hours after which the mesh cups containing the adult animals were removed, with the eggs falling through the mesh and remaining in the beaker. Another 24 hours later, hatched nauplii were counted and placed in groups of 20 in 16 beakers (4 replicates per station and salinity) filled with 200 mL of filtered sea water at the experimental and control salinity (7 and 15 PSU respectively). Previous experiments had shown high mortality at 5 PSU and therefore we chose 7 PSU as a low salinity condition in this experiment. In each beaker, *Rhodomonas* sp. was added at a concentration of 500 μgC/L and beakers were covered with aluminum foil to prevent evaporation. Survival was monitored after 3, 7, 10, 14 and 21 days by gently pouring the content of the beakers through a 50 μm sieve and transferring the animals to a petri dish to count them under a stereo microscope. Survival and developmental stages were noted, distinguishing the following categories: nauplius, copepodite stage C 1-3, C 4-5 and C6 (adults). For adult individuals the sex was noted. For the statistical analysis we used a generalized linear mixed effects model (GLMM) accounting for repeated measurements using *lme4* (Bates et al., 2015).

### Gene expression experiment

Because the cultures could be a mix of developmental stages, cultures were split to separate eggs from adults, and the eggs were used to start a new, synchronized culture where growth was monitored to catch the onset of maturation. For each population, 15 replicates of 50 newly matured adults (750 total) were counted and transferred into cups with a 100 μm mesh bottom for water exchange. The cups were inserted into three 20 L acclimation tanks at culturing conditions (15 PSU, 500 μgC/L *Rhodomonas*) and animals were allowed to recover from handling overnight. Before the start of the experiment, a control (t0) was sampled from each tank by swiftly transferring the experimental cups into RNAlater; for this and all following treatments there were three independent biological replicates. The remaining 12 cups were then placed into 7.5 L treatment tanks with a low salinity treatment (7 PSU) and a control treatment (15 PSU). Three hours post-transfer, animals were sampled for the first time point (t1). After 24 hours, all remaining animals were sampled (t2). Animals from the experimental cups were transferred to fresh RNAlater and split into two 1.5 mL tubes each containing 25 animals. Samples were stored at -20°C until processing.

### RNA extraction and sequencing

RNA was extracted from pooled animals (N = 25) using TRIzol reagent. For sample purification Qiagen RNeasy spin columns were used adding a DNAse treatment and second buffer RPE wash to improve sample purity. Sample quality and concentration was assessed using Qubit Fluorometer (Invitrogen) and Nanodrop (Thermo Fisher). Samples were stored at -80 °C before sequencing. For each biological triplicate, the subsample with the best quality was chosen for sequencing. Library preparation and sequencing was performed by Novogene UK on an Illumina Novaseq X.

Raw reads were quality assessed using *FastQC* (Andrews, 2010) and *MultiQC* (Ewels et al., 2016), adapters were trimmed using *fastp* (Chen et al., 2018). We then used *salmon* (Patro et al., 2017) to map reads to an existing *Acartia tonsa* transcriptome with ENA accession HAGX01, (Jørgensen et al., 2019). Mapping rate was 73.65 ± 0.37%. After filtering out low counts, 25,567 transcripts were included in the downstream analysis, 13,872 of which had a functional annotation.

### Transcriptomic analysis

The differential gene expression analysis was done in R v. 4.2.2. utilizing the *DESeq2* package (Love et al., 2014). We visualized variance stabilized transformed and regularized log transformed data in PCA plots highlighting time and treatment for both stations (Fig. S5). We employed two different approaches to analyze differential gene expression. First, we used a Likelihood-Ratio-Test (LRT) to extract transcripts that 1) were significantly differentially expressed at low salinity 2) showed a time-dependent expression pattern at low salinity 3) showed a population-dependent expression pattern at low salinity 4) showed a population and time dependent expression pattern at low salinity. Due to the imbalanced model design with no treatment applied at the control time point 0, it was necessary to manually edit the model matrices used as input for the LRT models in accordance with the *DESeq* manual. Second, to get a more detailed insight into genes that had a time-dependent salinity expression, we collapsed descriptors into an interaction term and used pairwise comparisons to contrast treatment and time point combinations for both stations separately. In all cases, transcripts were included in the final gene set if adjusted p-values were lower than 0.05 using the Benjamini-Hochberg false discovery rate as correction. Genes were considered overexpressed if log_2_fold changes were > 0 and underexpressed if log_2_fold changes were < 0. Differentially expressed genes were normalized and visualized using the *pheatmap* package (Kolde, 2019). To visualize general patterns of expression, we used the *degPatterns* function from *DEGreport* (Pantano, 2022).

To test for functional enrichment in the target gene sets, we used over representation analysis (ORA) and gene set enrichment analysis (GSEA) with gene ontology (GO) from *clusterProfiler* (Wu et al., 2021). Only genes that remained after filtering were included in the gene background against which the significant sets were tested for GO enrichment and an adjusted p-value cutoff of 0.05 was used (using the Benjamini-Hochberg false discovery rate correction). For the gene sets identified with pairwise comparisons, we used the *compareCluster* function of *clusterProfiler* and visualized the output with the *dotplot* function (Wu et al., 2021).

## Results

### Physiological assays

Physiological assays revealed both conserved and divergent plastic responses to acute salinity challenge. Salinity had an impact on survival for both populations, where survival was significantly lower at low salinities (p < 0.001) and decreased from 2 hours to 24 hours (p = 0.003; Fig. 1C). However, North Sea copepods had a lower survival at both timepoints (population effect; p = 0.012) and a significant salinity and population interaction indicated that Baltic Sea copepods were able to tolerate and survive in lower salinities to a greater degree than North Sea copepods (p = 0.004, LS_50_ after 2 hours: 0.76 ± 0.25 (Baltic), 1.10 ± 0.16 (North), and 24 hours:1.34 ± 0.52 (Baltic), 2.46 ± 0.19 (North)) (Fig. 1 B and C). Oxygen consumption of acclimated individuals did not differ in their response to salinity challenge. After overnight acclimation at 5 and 15 PSU, respiration rates were significantly different between stations (p < 0.001, Fig. S2), but not with treatment salinity (p = 0.844). Baltic organisms had higher oxygen consumption at both treatment and control compared to North Sea copepods: 5 PSU: 0.387 ± 0.132 (Baltic), 0.192 ± 0.064 (North), 15 PSU: 0.392 ± 0.079 (Baltic), 0.174 ± 0.046 (North), all measurements in μmol O_2_ h^-1^ mg dry body weight^-1^. In contrast, when challenged acutely, salinity had a significant effect on respiration rates (p < 0.001, Fig. 1D) but not station (p = 0.701) with respiration rates decreasing with decreasing salinity. Acutely challenged animals had higher respiration rates at all treatments, compared to the measurements after overnight acclimation: 5 PSU: 0.555 ± 0.104 (Baltic), 0.684 ± 0.194 (North), 2 PSU: 0.431 ± 0.065 (Baltic), 0.466 ± 0.141 (North). This observed effect of acclimation time on oxygen consumption can likely be explained by the higher metabolism of non-starved animals (Kiørboe et al., 1985).

Nauplii survival was also significantly influenced by treatment (p < 0.001) and time (p < 0.001). There was a significant effect of population on survival with Baltic copepods showing higher survival (p = 0.013). However, the low treatment salinity of 7 PSU was lethal for nauplii from both populations where no individual reached the copepodite stages and no individuals survived beyond 14 days (Fig. S3). At the control salinity, adults were recorded after 14 days with all surviving individuals reaching adulthood after 21 days (North Sea: 42.5 % ± 6.5 %, Baltic Sea: 50.0 % ± 10.0 %). The female:male ratio differed drastically between populations (2.08 for Baltic Sea, 0.28 for North Sea). However, this observation of sex ratios does not reflect our usual experiences from handling these cultures where anecdotally sexes are relatively even, and is potentially an artifact of picking a subset of eggs.

Lastly, neither egg production nor hatching were significantly influenced by salinity (p = 0.737, and p = 0.517 respectively) or population (p = 0.281, and p = 0.658 respectively). Daily egg production was highly variable and averaged at 34.3 ± 25.8 for the North Sea population and 26.8 ± 14.0 for the Baltic population across all treatments. For the North Sea population 20.8 ± 15.9 nauplii hatched (60.7 % of all eggs) and for the Baltic Sea population 18.8 ± 10.3 nauplii hatched (70.3 % of all eggs, Fig. S4).

### Gene expression

We first identified genes that showed a significant response to low salinity using the LRT model. We found 889 differentially expressed genes (DEGs) for which gene ontology (GO) enrichment yielded 8 terms that were significantly enriched in both ORA and GSEA. Here, the biological processes were exclusively related to transport functions (GO:0055085 transmembrane transport, GO:0006811 ion transport, GO:0006814 sodium ion transport, see Table S1). Of these DEGs, 105 were significant for an interaction between low salinity and time, with enrichment for 18 GO terms, 7 of which were biological processes. Again, functional enrichment of biological processes was dominated by transport processes (GO:0055085 transmembrane transport, GO:0006811 ion transport, GO:0006814 sodium ion transport, GO:0006869 lipid transport, see Table S2). Though the LRT only identified two genes as differentially expressed for the two-way interaction between low salinity and population, and one for the three-way interaction between low salinity, population and time, the PCA of the 889 significant salinity transcripts shows clear clustering by both time and population (Fig. 2). We therefore opted for pairwise comparisons to gain further insights into more subtle population differences in contrast to the broad patterns identified by the LRT (Love et al., 2014).

**Figure 2.**
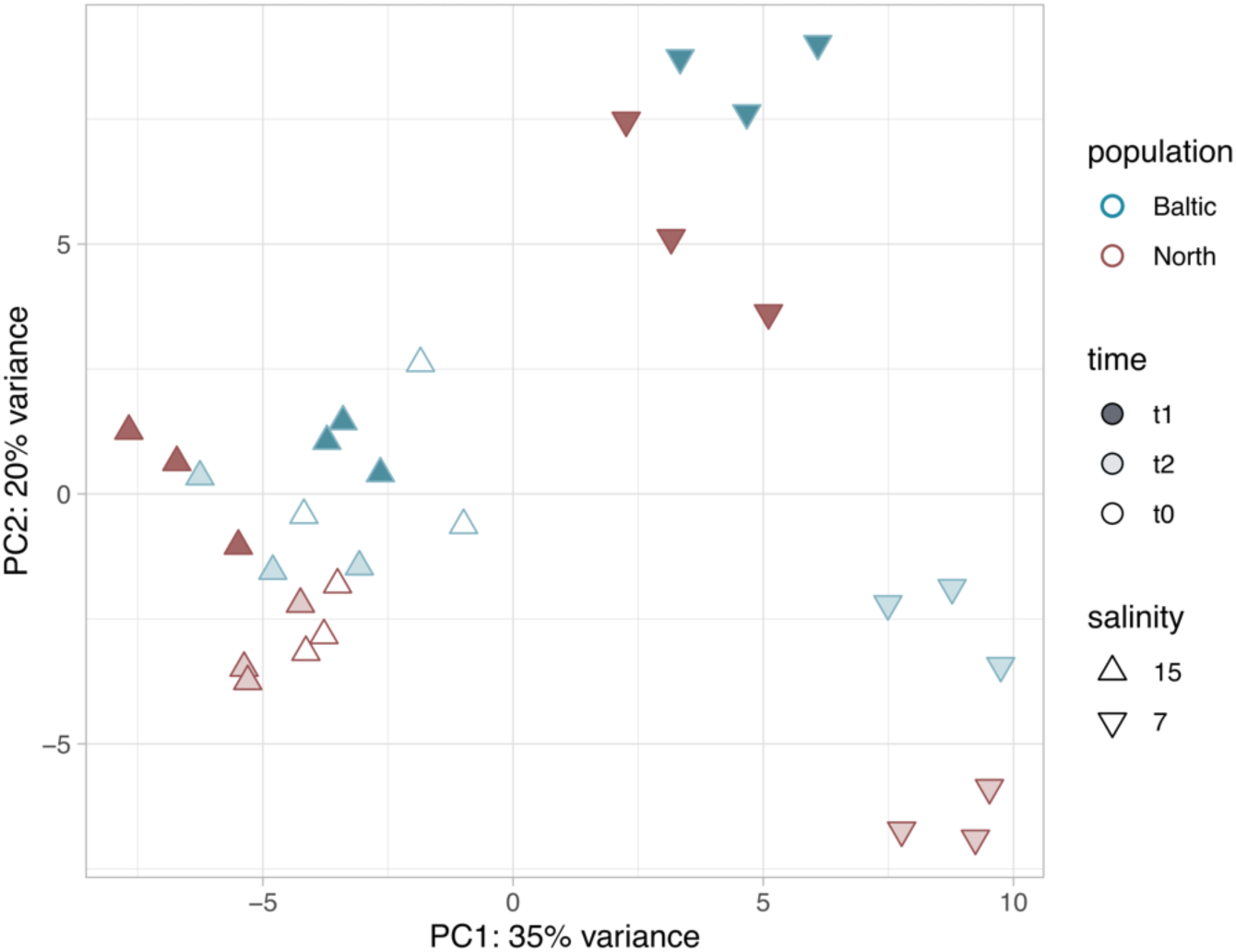
PCA of 889 genes significantly differentially expressed at low salinity (Likelihood Ratio Test). Samples from the Baltic are shown as blue triangles and the North Sea as red triangles. Treatment salinity was measured in practical salinity units (PSU), t0 is the control sampling before transfer to the treatment salinities, t1 and t2 were sampled 3 and 24 hours post transfer, respectively.

When using pairwise comparisons, contrasting the low salinity treatment for each station and time point with the respective controls, 341 DEGs were identified at low salinity for either t1 or t2 in the Baltic population, and 415 DEGs for the North Sea copepods, totaling to 602 unique salinity-dependent genes identified by pairwise comparisons. Of these, 516 DEGs were shared with the salinity-dependent LRT (85.7% of all pairwise transcripts, and 58.0% of all LRT transcripts).

### Conserved salinity response

Across both populations, 154 pairwise genes were jointly differentially expressed at low salinity (Fig. 3), of which 63 were also identified by the time-and salinity-dependent LRT. This conserved plastic response to salinity included 79 transcripts for the short-term response (t1, 3h) and 91 transcripts for the long-term response (t2, 24 h) highlighting time-dependent shifts in gene regulation.

**Figure 3.**
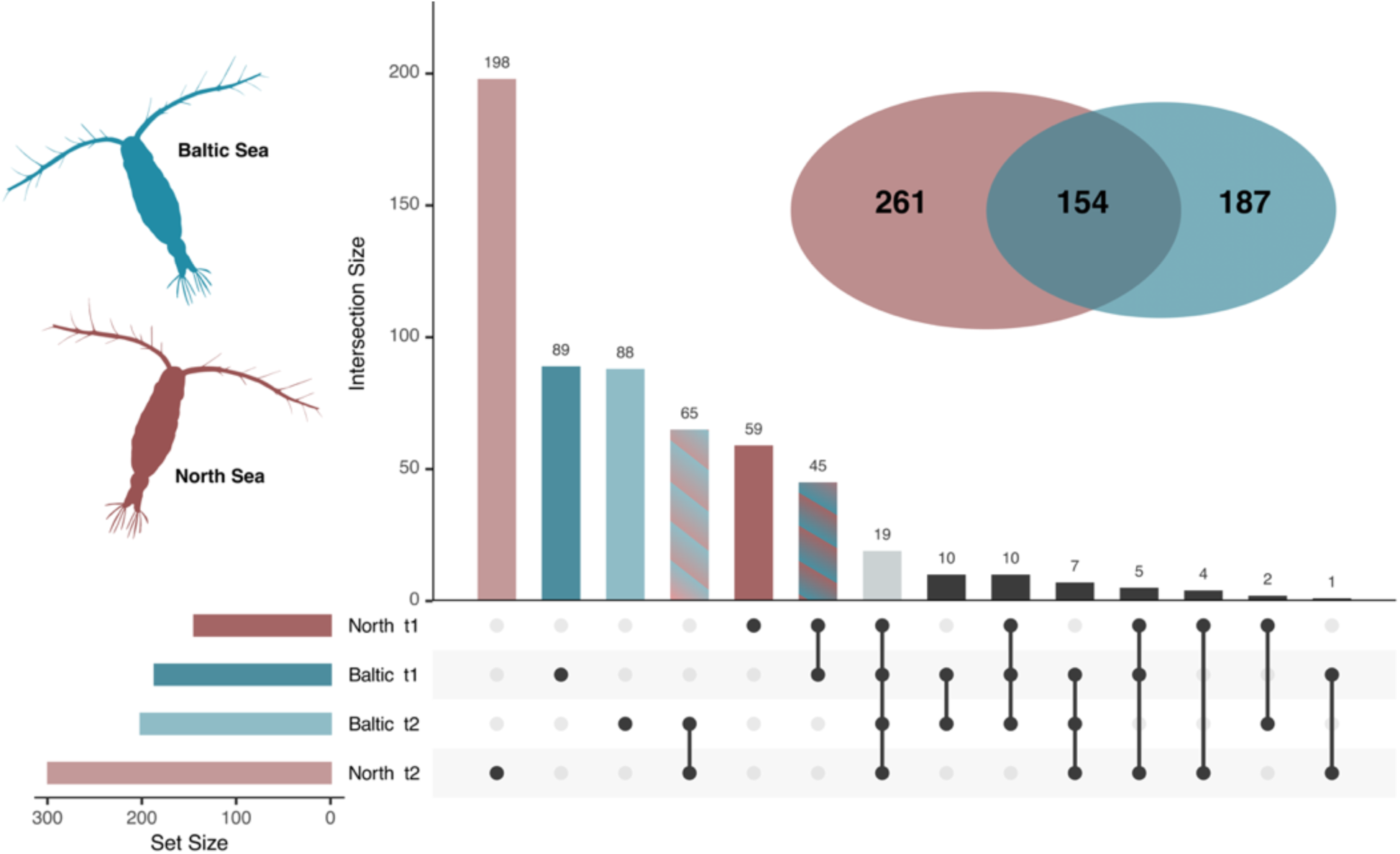
Counts of significantly expressed genes for Baltic (labeled in blue) and North Sea (labeled in red) *Acartia tonsa* exposed to low-salinity stress for 3 hours (t1) and 24 hours (t2) identified with pairwise comparisons. The Venn diagram shows the overall significant genes at both time points as well as the overlap between populations. The upset plot shows unique and intersecting genes between populations and time.

GO enrichment revealed time-dependent patterns of biological processes in response to low salinity exposure. In the short-term response, functional enrichment was primarily driven by upregulated genes, with key processes related to osmoregulation and ion transport (GO:0055085 transmembrane transport, GO:0015701 bicarbonate transport, GO:0006820 anion transport). Additionally, metabolic and neurological regulation was enriched, including GABA metabolism (GO:0009450 and GO:0009448) and glucose metabolic process (GO:0006006). Other enriched functions suggested a systemic response to salinity stress, such as such as positive regulation of heat generation (GO:0031652), modulation of inhibitory postsynaptic potential (GO:0097151), and behavioral responses (GO:0048148 behavioral response to cocaine) (Fig. 4, full list in Table S3). Several DEGs associated with osmoregulatory processes were identified, including sodium-dependent glucose transporters (*G5CMC2* and *RPPX_22905*), a major facilitator superfamily (MFS) type transporter (*SLC18B1*), and carbonic anhydrase (*CAA6C*) suggesting active ion regulation (full list in Table S4).

**Figure 4.**
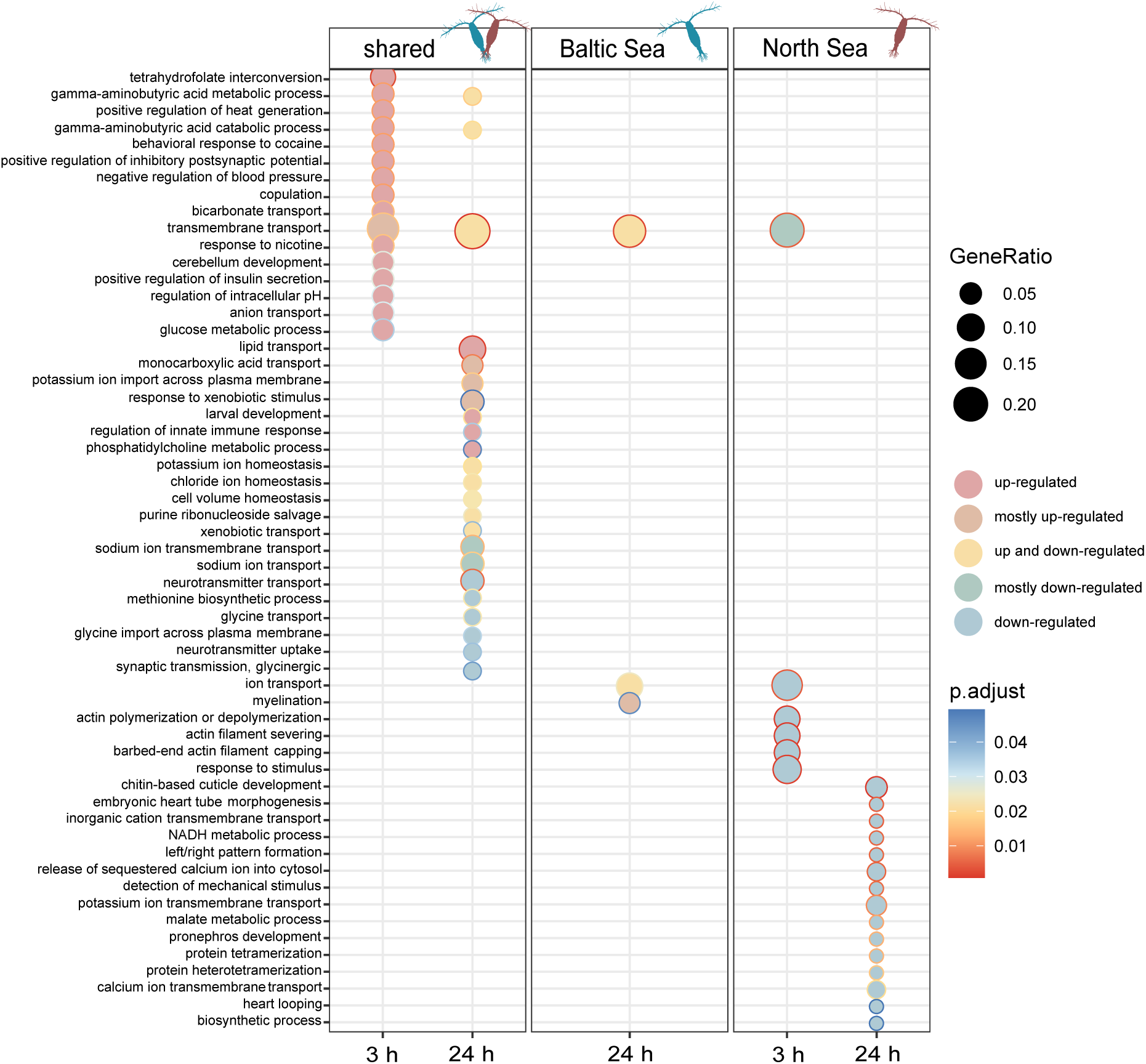
Gene ontology (GO) enrichment for biological processes (BP) of time-dependent salinity genes for shared, and unique genes of Baltic and North Sea *Acartia tonsa.* Enrichment was calculated with the *compareCluster* function of *clusterProfiler* (v 4.0, Wu et al., 2021). Regulation of genes within each GO term is indicated by fill; outline color indicates adjusted p-values (using the Benjamini-Hochberg false discovery rate correction).

After 24 hours of exposure, transmembrane transport remained a key enriched function, but transport-related processes diversified, with both upregulated and downregulated transport pathways emerging. Upregulated genes were associated with lipid transport (GO:0006869), monocarboxylate transport (GO:0015718), and potassium ion import (GO:1990573), while downregulated genes drove enrichment in sodium ion transmembrane transport (GO:0035725) and glycine transport (GO:0015816). The prolonged stress response was also characterized by enrichment in stress response mechanisms, including response to xenobiotic stimulus (GO:0009410) and regulation of innate immune response (GO:0045088). Additionally, key homeostatic processes were enriched, such as potassium ion homeostasis (GO:0055075), chloride ion homeostasis (GO:0055064), and cell volume homeostasis (GO:0006884), indicating a shift towards long-term osmoregulatory adjustments. Notably, genes associated with these enriched processes included a solute carrier (*SLC12A3*), an organic cation transporter (*CGI_10003241*), and a carbonic anhydrase (*CA4-like*), further supporting the role of active ion regulation in coping with salinity stress.

### Population specific response

A total of 187 and 261 genes showed unique differential expression in response to low salinity for the Baltic and North Sea populations, respectively (Fig. 3). For the Baltic population, 99 DEGs (89 unique to t1; 10 shared between t1 and t2) were differentially expressed during short-term exposure (t1), while 98 DEGs (88 unique to t2; 10 shared between t1 and t2) were differentially expressed after 24 hours (t2). No biological functions were significantly enriched at t1. However, by t2, upregulated genes predominantly contributed to enrichment in three key biological processes: Transmembrane transport (GO:0055085), ion transport (GO:0006811) and myelination (GO:0042552) (Fig. 4). Among the differentially expressed osmoregulatory genes, cation transporting ATPases (*ATP13A3* present at both time points), organic cation transporters (*R4UKP6, OCT1*), and an MFS transporter (*Orct_13*) were identified.

In contrast, the North Sea population exhibited a more extensive transcriptional response over time with 63 DEGs (59 unique to t1; 4 shared at t1 and t2) identified for the short-term response, increasing to 202 DEGs (198 unique to t2 + 4 shared at t1 and t2) at t2. At both time points, functional enrichment was driven by downregulated genes, suggesting a suppression of key biological processes. At t1, downregulated functions were associated with structural reorganization, including actin filament severing (GO:0051014) and actin filament capping (GO:0051016), alongside suppressed ion and transmembrane transport processes (GO:0006811, GO:0055085) (Fig. 4). Notably, osmoregulatory genes downregulated at this stage included receptor potential cation channels (*TRPA-1*). By t2, suppression extended beyond transport processes to include biosynthesis and metabolism (GO:0009058 biosynthetic process, GO:0006108 malate metabolic process, GO:0006734 NADH metabolic process), developmental pathways (GO:0048793 pronephros development, GO:0008362 chitin-based cuticle development), and ion homeostasis mechanisms (GO:0098662 inorganic cation transmembrane transport, GO:0071805 potassium ion transmembrane transport, GO:0070588 calcium ion transmembrane transport). Among the significantly downregulated osmoregulatory genes at this stage were voltage-gated potassium channels (*KCNV1*, *AMK59_7147*).

When comparing relative expression patterns, genes differentially expressed at low salinity generally exhibited higher expression levels in the Baltic population (Fig. 5). While the overall direction of gene expression changes was consistent between populations, the North Sea population displayed relatively lower expression levels at low salinity, particularly after 24 hours (Fig. S6).

**Figure 5.**
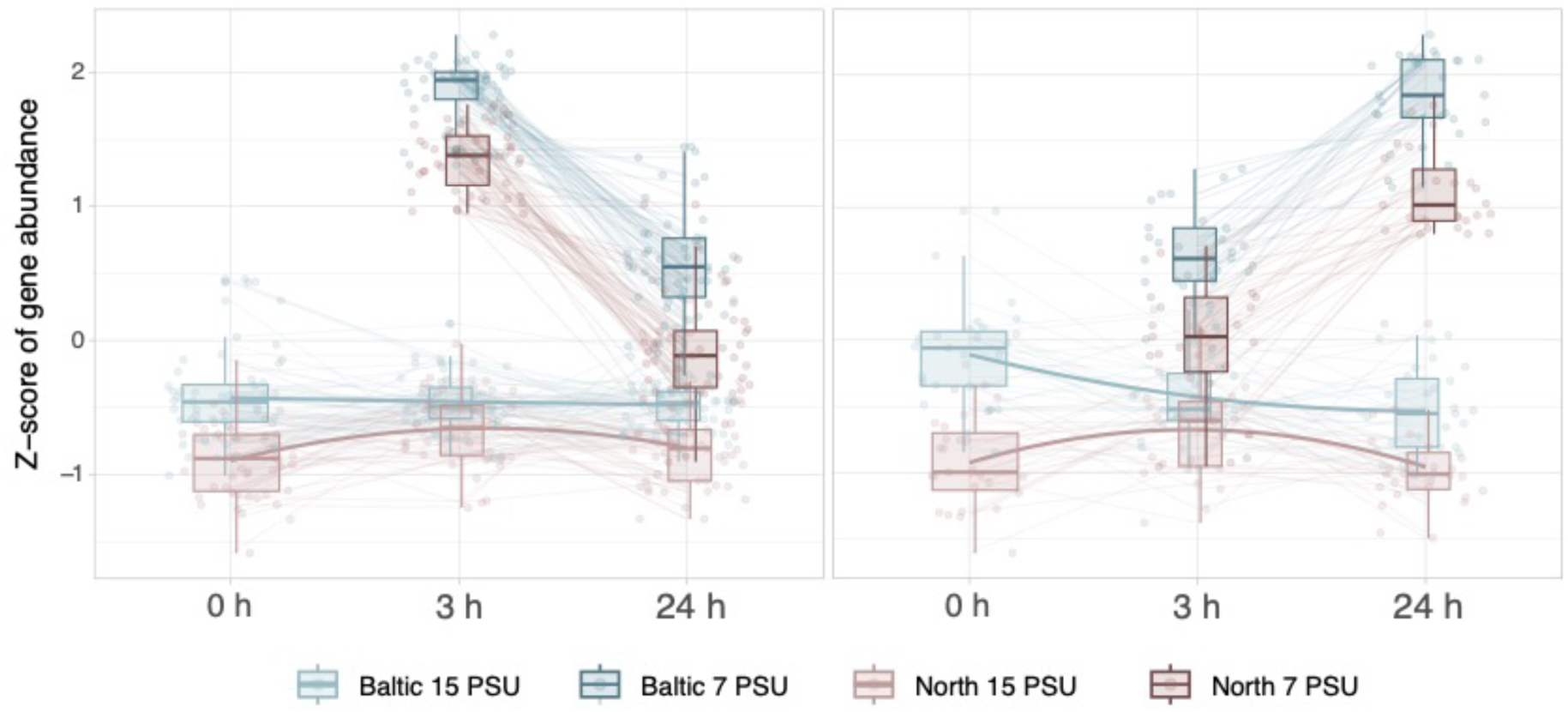
Gene clusters with similar expression patterns at low salinity show population differences in expression levels. Left: 48 of 154 shared genes differentially expressed at low salinity treatment; Right: 27 of 154 shared genes differentially expressed at low salinity treatment. Clustering performed using *degPatterns* from *DEGreport* (Pantano, 2022).

## Discussion

We used physiological experiments on copepods common-gardened for multiple generations combined with a transcriptomic assay to investigate local adaptation to salinity in a Baltic and North Sea population. If local adaptation to low salinity was present, we expected to see higher fitness in the low salinity Baltic Sea copepods than those originating from the high salinity North Sea when exposed to low salinity stress. The increased survival of the Baltic Sea individuals at low salinity relative to the North Sea population is indicative of local adaptation to salinity. However, not all phenotypic measures supported this conclusion as there were no consistent differences in metabolism and reproduction. The gene expression responses allowed us to identify the conserved plastic response to low salinity stress that enables *A. tonsa* as a species to tolerate variable osmotic environments and, as might be expected, included functions related to ion transport and homeostasis (Fig. 4). Population level differences in expression were primarily driven by increased differential expression in the high salinity North Sea population, suggesting an inability to efficiently osmoregulate and tolerate low salinity conditions relative to the Baltic Sea population. Our findings suggest local adaptation to the low salinity conditions in the Baltic Sea with an expansion of plasticity to tolerate these dilute conditions but also highlight the extraordinary plasticity of *A. tonsa* as a species across broad osmotic environments.

### Phenotypic traits reveal local adaptation

Baltic Sea and North Sea populations showed phenotypic differences in response to low salinity challenges that were consistent with local adaptation. As expected, extreme low salinity increased mortality in both populations, but Baltic Sea copepods showed significantly higher survival than their North Sea counterparts. Despite this evidence for local adaptation in short-term survival, other physiological and reproductive traits did not differ between populations. Oxygen consumption rates were similarly affected by low salinity in both populations, and reproductive success - measured by egg production and hatching rates - remained unchanged.

Typically, *A. tonsa* exhibits an increase in respiration rates when exposed to salinities deviating from its native environment (Gaudy et al., 2000; Lance, 1965), but salinities below 5 PSU have not been considered in past experiments. In extremely low salinity environments copepods reduce their swimming activity (Seuront, 2006), which we anecdotally observed in our low salinity treatments. This behavioral response could explain why, despite experiencing osmotic stress, overall metabolic rates were reduced.

Our experiments simulated an acute salinity shock, mirroring natural salinity reductions during heavy rain falls or tidal cycles. In these cases, evolving a short-term strategy to tolerate extreme hyposaline conditions is necessary to ensure survival. Prolonged exposure could be avoided by behavioral responses such as migrating to deeper, more saline layers. Indeed, decreased surface salinity alters the vertical migration patterns of copepods (Lance, 1962) likely to avoid prolonged exposure to harmful conditions. As a consequence, our experimental setup enabled us to uncover population differences in survival but not reproductive success since the latter trait might not be impacted by short-term exposure of adult individuals to low salinity. This is especially true since parental investments in egg production likely have been made prior to the start of the experiments (Niehoff, 2007). In similar experiments on the reproduction of *Acartia* with longer acclimation periods, egg production rate decreased at low salinities (Dutz & Christensen, 2018; Peck & Holste, 2006).

While short-term responses appear critical for immediate survival, the long-term strategy to tolerating the overall reduction in salinity in the Baltic Sea may include more subtle phenotypes that would only emerge under long-term acclimation or development at low salinities. We initially planned to assess reproductive output in copepods reared at low salinity, but complete naupliar mortality at these conditions (Fig. S2) prevented further experiments. This suggests that, at least in the populations used in this study, there may be a lower limit to population persistence under salinity decreases and that developmental stages differ in their salinity tolerance (Magouz et al., 2021; Nour et al., 2021). Future studies should quantify the long-term effects of low salinity exposure on fitness in *A. tonsa* from a suite of osmotic environments.

### Core mechanisms of osmoregulatory plasticity in *A. tonsa*

Typically, most invertebrates are isoosmotic with equal osmolarity to their surrounding and a decrease of salinity leads to an inflow of water into the cells that is initially counteracted by the release of inorganic ions such as K^+^, which reduces internal osmolarity and helps prevent further swelling (Moran & Pierce, 1984). Since ion concentrations are important for cell functions due to their role in maintaining membrane potentials, over intermediate timeframes compatible organic osmolytes such as free amino acids (FAA) are used to maintain isoosmotic conditions (Otto & Pierce, 1981; Yancey, 2005). When stressful conditions persist for longer time periods or the salinity reaches a critical lower limit, organisms are forced to engage in energetically costly active ion regulation to sustain cell function (Lee et al., 2011), and as consequence organisms become hyperosmotic compared to their surroundings (Mantel & Farmer, 1983).

The osmoregulatory strategy of *A. tonsa* previously has not been extensively investigated. *Acartia tonsa* has been found to be a weak hyperosmotic osmoregulator, maintaining isoosmotic body fluids at native salinities and becoming slightly hyperosmotic when salinity decreases (Lance, 1965; Mantel & Farmer, 1983). There is evidence for osmoregulatory efforts such as regulation of ion concentrations and amino acid levels to maintain constant cell volumes in low salinity (Farmer & Reeve, 1978; Svetlichny & Hubareva, 2014). This is consistent with the osmoregulatory mechanism of other copepods. For example, both the cyclopoid copepod *Apocyclops royi* (Jepsen et al., 2025) and *Eurytemora affinis* are isoosmotic at higher salinities, but osmoregulate when exposed to low salinity (Gerber et al., 2016; Johnson et al., 2014).

Our findings indicate that *A. tonsa* likely follows the standard model of invertebrate osmoregulation, showing both short term regulation of ions as well as an intermediate regulation of amino acids to maintain homeostasis. The large core set of genes that consistently responded to the low salinity challenge between Baltic and North Sea populations illuminates how *A. tonsa* is able to tolerate considerable salinity decreases.

Under low salinity stress, it is necessary to control both intercellular ion homeostasis and cell volume which require active transmembrane transport and water balance, which are energetically costly and can affect intracellular pH levels (Krasznai et al., 1995; Kültz, 2015). Accordingly, the immediate response to low salinity was dominated by metabolic upregulation, as well as ion transport and pH regulation in an initial effort to combat salt stress and maintain homeostasis.

With prolonged exposure to low salinity, the functional response broadened beyond ion transport to include the transport of larger molecules such as amino acids (mainly glycine) and lipids, indicating a shift in osmoregulatory mechanisms. Glycine, an important organic osmolyte in many marine invertebrates, plays a key role in maintaining cellular osmotic balance under hypo-osmotic stress (Podbielski et al., 2022). The regulation of lipid transport may support plasma membrane integrity during persistent osmotic stress (Mu et al., 2024). The activation of diverse transport functions in response to low-salinity stress has been observed across multiple marine taxa, including euryhaline fish, oysters, and copepods (DeBiasse et al., 2018; Jones et al., 2019; Li et al., 2025).

In many crustaceans, Na^+^/K^+^-ATPase and V,H^+^-ATPase (VHA) are important osmoregulatory genes (Gerber et al., 2016; Lee et al., 2011; Lv et al., 2013; Posavi et al., 2020) and we expected to observe these candidate genes in the core response to salinity in *A. tonsa*. While these specific genes were not differentially expressed in our samples, we found a suite of ion transporters responding to salinity challenges, including Na^+^ and Cl^-^ dependent transporters. It is likely that these transporters are serving a similar role in contributing to osmoregulation. For example, at t1 and t2 both populations showed upregulation of the sodium chloride cotransporter (*NCC*; *SLC12A3*), that reabsorbs Na^+^ and Cl^--^ in the urinary bladder of teleost fishes exposed to freshwater (Takvam et al., 2021). Alternatively, fully assembled ATPases could be stored in vesicles and be activated and translocated upon stress as has been shown with VHA in sharks (Tresguerres et al., 2010). This would allow animals to quickly respond to salinity changes without activating transcriptional pathways.

Further, we found *carbonic anhydrases* which are linked to osmoregulation in crustaceans and previously were shown to be upregulated under low salinity stress in the Pacific white shrimp (Roy et al., 2007). Carbonic anhydrases convert respiratory CO_2_ respiration to bicarbonate and protons to take up ions such as Na^+^ and Cl^-^ via e.g., anion exchangers (AE), VHA and sodium proton exchangers (NHE); carbonic anhydrases are likely core to the copepod osmoregulatory strategy (Lee, 2023; Lee, Charmantier, et al., 2022). In addition, the regulation of GABA pathways plays an important role, as this neurotransmitter can increase tolerance to abiotic stress by maintaining membrane potential and promoting ion transport and has been found to accumulate in response to salt stress in plants (Cheng et al., 2018; Su et al., 2019; Yuan et al., 2023). Further, *major superfacilitator family* (MFS) genes were shared between populations. MFS genes are a family of transporters operating under a chemiosmotic ion gradient and include sugar transporters (Pao et al., 1998) such as the *sodium-dependent glucose transporter 1* that was upregulated at t1. Finally, De Vos et al. (2019) identified several groups of transport genes that were differentially expressed in *Artemia* brine shrimp following salt stress, such as ion transporters and salt-dependent transporters. Similarly, we found various solute carrier genes including *organic cation transporter-like* or *organic anion transporter family member 2A 1*. Many of the genes or functional equivalents that we identified here were also differentially expressed in the copepod *E. affinis* under salinity stress (Posavi et al., 2020), indicating that they are likely important in osmoregulation across diverse copepods.

The time-sensitive pattern of the stress response we observed suggests a dynamic osmoregulatory strategy common to invertebrates. Initially, *A. tonsa* seems to mainly rely on active ion transport to counteract the immediate osmotic shock, regulating intracellular ion concentrations and pH levels. However, after prolonged exposure, the shift towards organic osmolytes and lipids may indicate a transition to a more stable state, potentially reducing the high energetic demands of active osmoregulation and avoiding depletion of essential ions. Future studies should investigate if *A. tonsa* can reach a fully isoosmotic state at the salinity we tested or if continued exposure to low salinity will force the organism to maintain hyperosmotic levels through active osmoregulation.

### Population specific gene expression underlying local adaptation

We identified three main differences in gene expression that may underlie local adaptation between the populations. First, more genes with a time-dependent salinity response were expressed in the North Sea population. From 3 hours to 24 hours this population more than tripled the number of population-specific DEGs while the number of population-specific DEGs in the Baltic Sea population remained stable (Fig 3). The lower numbers of differentially expressed genes may reflect increased resilience of the Baltic Sea population to low salinity stress as they are able to recover and maintain homeostasis more efficiently, similar to transcriptomic resilience (Franssen et al., 2011). The European flounder (Larsen et al., 2007) and the copepod *Tigriopus californicus* (DeBiasse et al., 2018) similarly have muted transcriptomic responses that correspond to higher salinity tolerance, plasticity, and local adaptation. In addition, the differentially expressed genes were almost exclusively downregulated, meaning that functions were being suppressed, suggesting that the North Sea copepods were prioritizing survival over growth-limiting metabolic demands and costly active ion transport especially after prolonged exposure to low salinity (Hand & Hardewig, 1996).

Second, many of the genes associated with low salinity responses in both populations were more highly expressed in the Baltic population during the stressful condition, and in some cases also the control state. This is similar to the phenomenon known as ‘frontloading’ that has been described first in corals where resilient corals showed higher expression levels of important stress related genes in an undisturbed state (Barshis et al., 2013). We suggest that the Baltic population could have adapted to low salinity conditions by evolving an increased basal expression of key osmoregulatory genes, allowing them to more efficiently respond to sudden decreases in salinity.

Lastly, some genes which have been linked to be osmoregulation in crustaceans were uniquely differentially expressed only in a single population. North Sea copepods showed differential expression of *arginine kinase,* an enzyme that facilitates ATP formation by transferring phosphate from phospo-L-arginine, the phosphagen ATP buffer in many invertebrates (Holt & Kinsey, 2002). In the blue crab *Callinectes sapidus,* hypo-osmotic conditions induced a higher arginine kinase flux and in the estuarine copepod *E. affinis*, arginine kinase was identified as key osmoregulatory gene in response to fresh water exposure (Holt & Kinsey, 2002; Posavi et al., 2020). Only Baltic copepods showed significant regulation of *ATPase* genes (e.g., *ATP13A3, P-type ATPase*) which use ATP as an energy source to fuel active exchange of ions across the cell membrane. These differences in expression of osmoregulatory genes could point towards adaptation of the Baltic population to the decreased salt levels in their natural habitat and explain the higher survival we observed in this study.

### Conclusions and implications for future changes

We showed that *A. tonsa* as a species is able to withstand extreme salinity conditions for hours to days. Given these findings, we assume that *A. tonsa* will be able to tolerate climate-driven salinity declines in most of its distribution, the exception being the north-east Baltic Sea. Since this low salinity area represents the outermost limit of the current species’ distribution, further decreases in surface salinity might lead to unfavorable conditions for larval development and therefore range contractions.

Despite their broad osmotic plasticity, we found evidence for local adaptation to low salinity, indicating that *A. tonsa* from higher salinity regions need to adapt to thrive under decreased salinity regimes. The Baltic Sea as we know it today is a new body of water, only about 8,000 years old (Björck, 1995), and the relatively rapid adaptation to low salinity observed here has been driven by an expansion of plasticity to tolerate the newly occupied habitat, consistent with other organisms moving from high to low salinity (Brennan et al., 2015; Lee et al., 2011). This expansion appears to be driven by a more efficient physiological response (Fig. 2) that includes front-loading of key osmoregulatory genes (Fig. 5).

The evidence for local adaptation presented here indicates that *A. tonsa* has the capacity to evolve tolerance to low salinities and – in the past – has adapted to novel environments. This suggests that the species potentially can adapt to future changes in ocean salinity following the space-for-time concept (Kharouba & Williams, 2024). Its high genetic variation and rapid generation times have enabled it to adapt to other contemporary anthropogenic changes over just tens of generations (Brennan, et al., 2022a), and this adaptive capacity could also extend to future changes in salinity. However, rapid evolution can have tradeoffs such as loss of plasticity or reduced fecundity (Brennan et al., 2022b). Indeed, habitat shifts and local adaptation from high to low salinity in other organisms have been accompanied by expansions of plasticity to tolerate low salinity, but a loss of plasticity to tolerate the ancestral environment (DeFaveri & Merila, 2014; Marchinko & Schluter, 2007) consistent with theoretical predictions of genetic assimilation (Lande, 2009). If *A. tonsa* originating from low salinity carry the plasticity to tolerate both low and high salinity fluctuations is currently not known, but adaptation to low salinity could carry costs. Future studies should address the limits of *A. tonsa’*s adaptive potential, the impacts of rapid adaptation on plasticity and investigate potential tradeoffs of rapid adaptation to low salinity.

## Acknowledgements

We thank Diana Gill, Fabian Wendt, and Claas Hiebenthal for their technical support in the lab and climate chambers, and Frank Melzner, Jasmin Renz, Thorsten Reusch, and Meike Stumpp for their scientific counsel. We are grateful for the crew and scientists aboard the AL580 cruise on RV Alkor (chief scientist: Felix Mittermayer-Schmittmann). Special thanks to Sheena Chung for taking part in the transcriptomic sampling event. This research was supported in part through high-performance computing resources available at the Kiel University Computing Centre. The project was funded by the Deutsche Forschungsgemeinschaft (DFG, German Research Foundation) - project number 504958481 to R.S.B.

## Data availability

The code and physiological data that support the findings of this study are archived and openly available at Zenodo (http://doi.org/10.5281/zenodo.15535306). Transcriptomic read data and sample metadata is available at NCBI under the project accession PRJNA1258960.

## Notes

### Competing Interest Statement

The authors have declared no competing interest.

