## Supplemental Figures for "Local Adaptation and Transcriptomic Plasticity of the Copepod *Acartia tonsa* Under Low Salinity Stress"

### Supplementary Figures

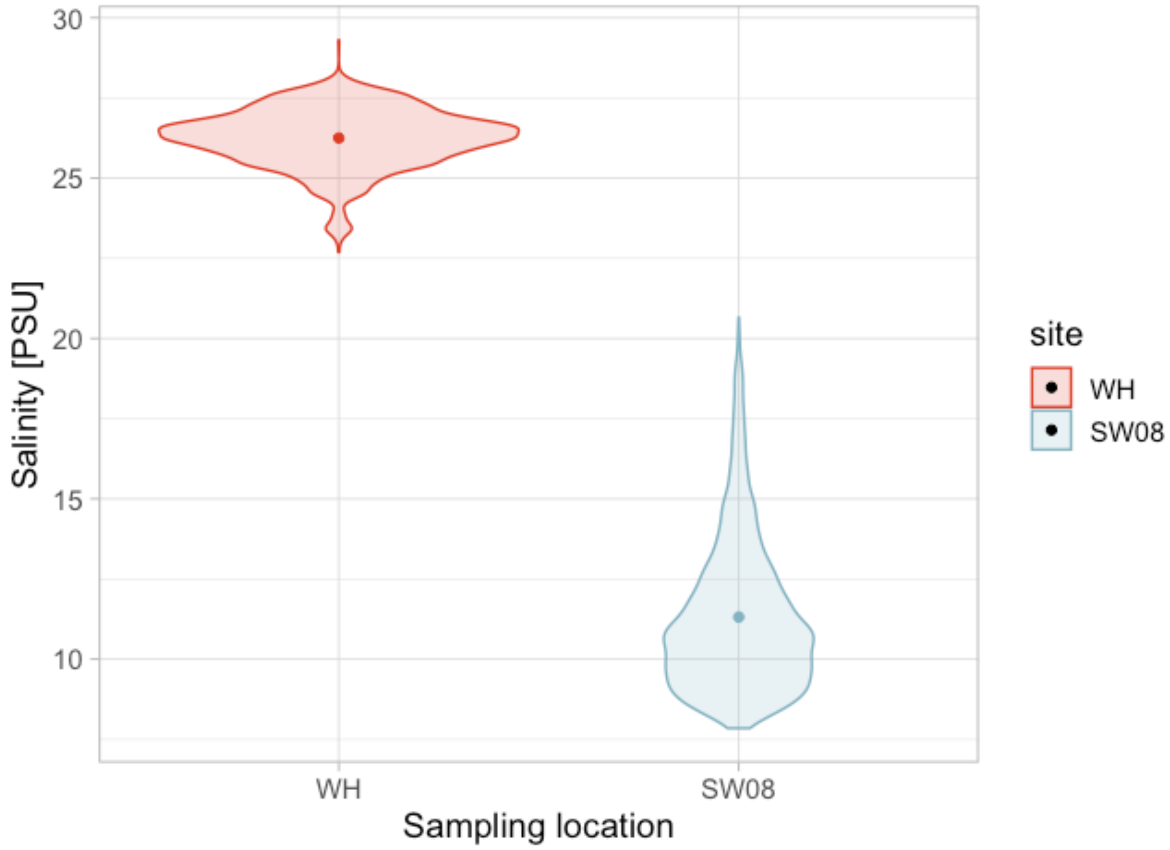


Figure S1. Daily surface salinity measurements for both sampling locations from 1993-2021, includes all measurements from June to October. Colored points indicate mean salinity values for the Baltic location (blue) and North Sea location (red). Data from CMEMS https://doi.org/10.48670/moi-00021

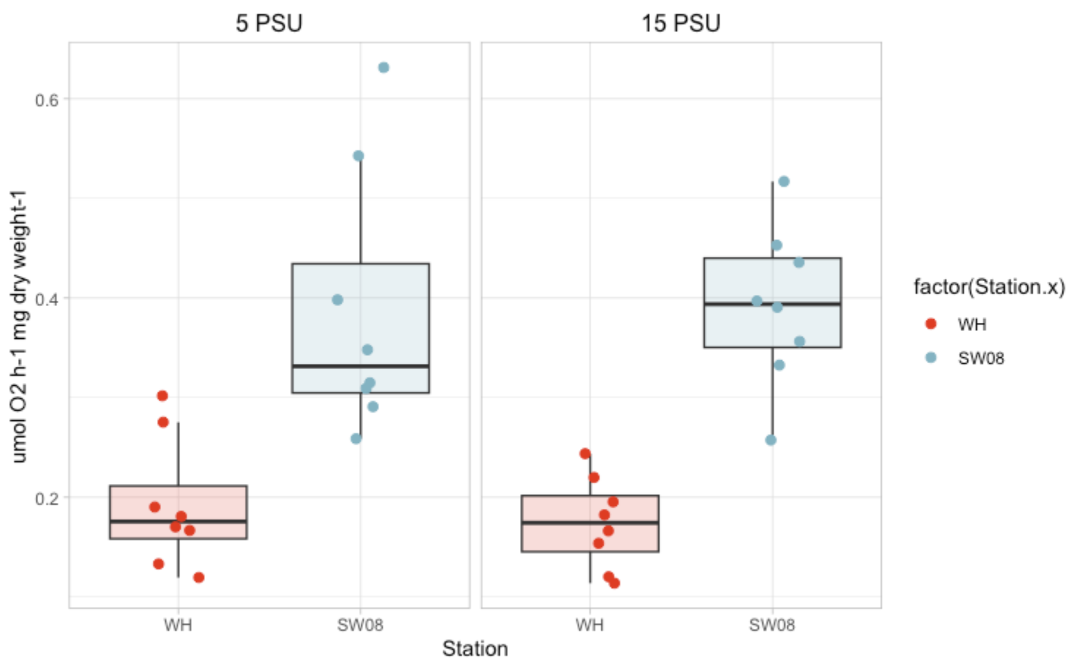


Figure S2. Respiration rates of *Acartia tonsa* individuals from the Baltic Sea (blue) and the North Sea (red), standardized per dry body weight. Measurements were conducted after 16 hours of acclimation to the treatment salinities.

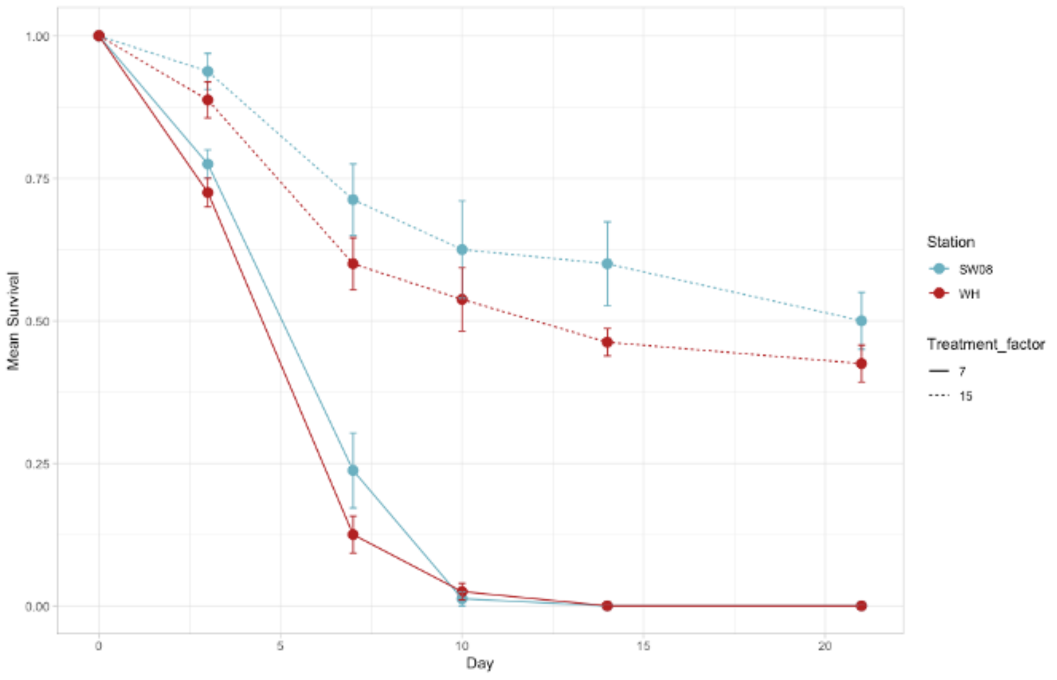


Figure S3. Naupliar survival of Acartia tonsa from the Baltic (blue) and North Sea( red) at two treatment salinities (7 and 15 PSU), points indicate mean survival at sampling days, error bars indicate standard deviations.

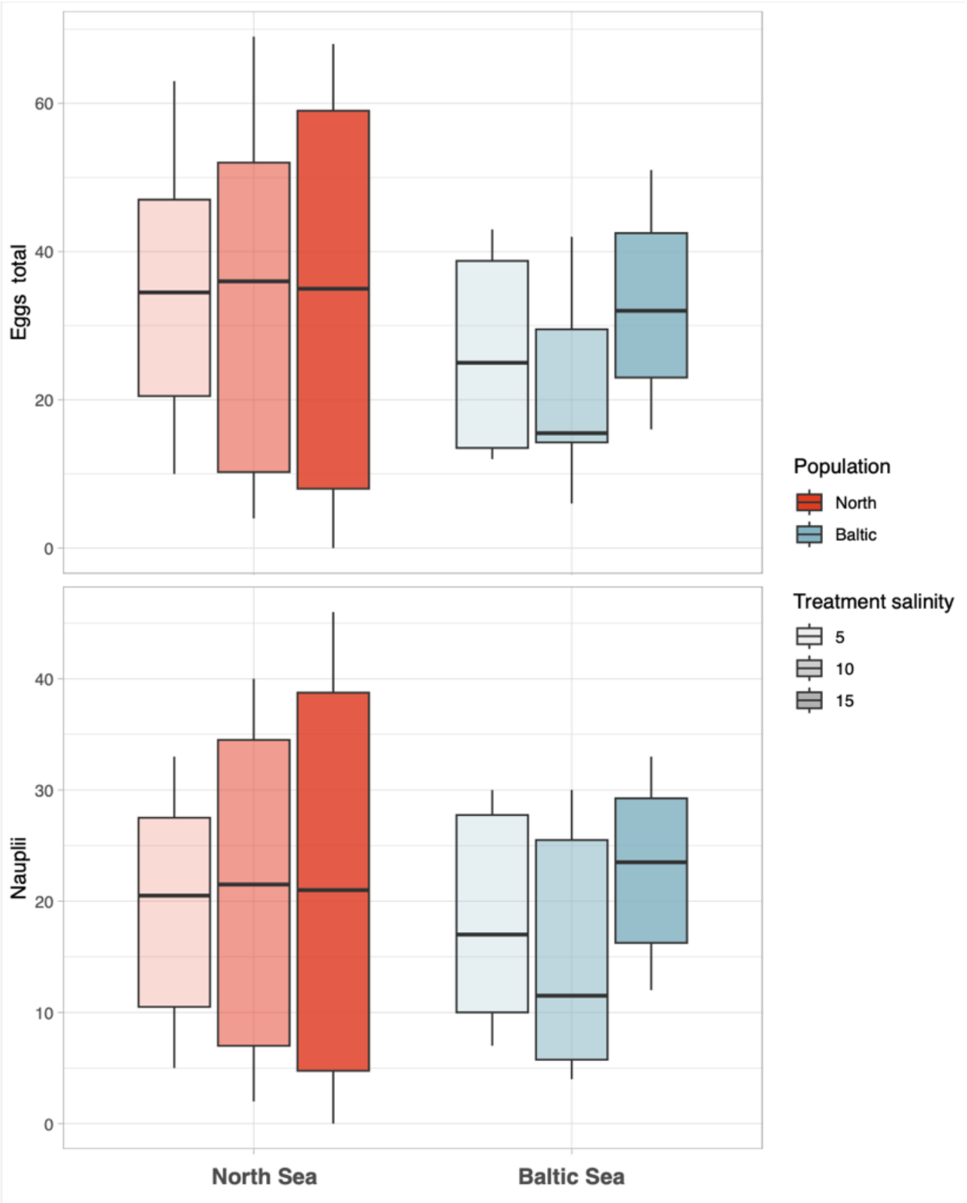


Figure S4. Egg production and hatching success of Acartia tonsa from the Baltic (blue) and North Sea (red) after acute transfer to three treatment salinities (5, 10, 15 PSU).

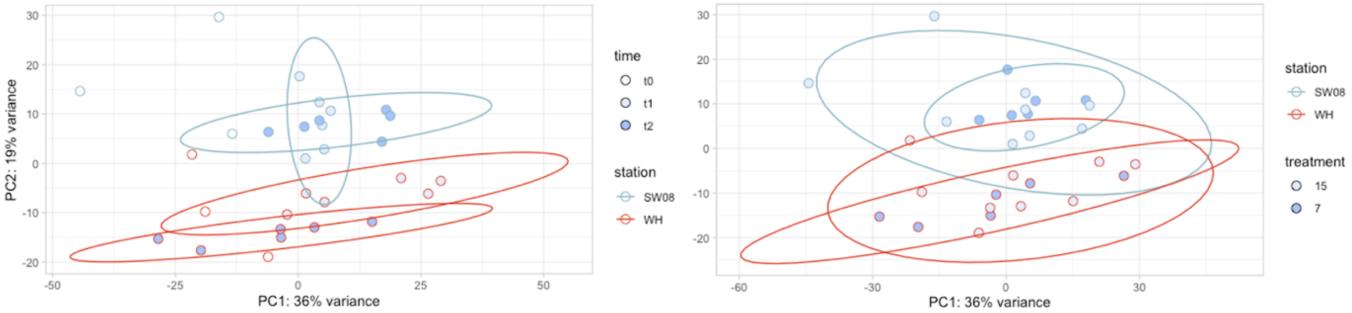


Figure S5. PCA of all 25,567 genes that remained passed filtering, the Baltic population is colored in blue, the North Sea population in red. Left: ellipses show clustering by population and time; Right: ellipses show clustering by population and treatment.

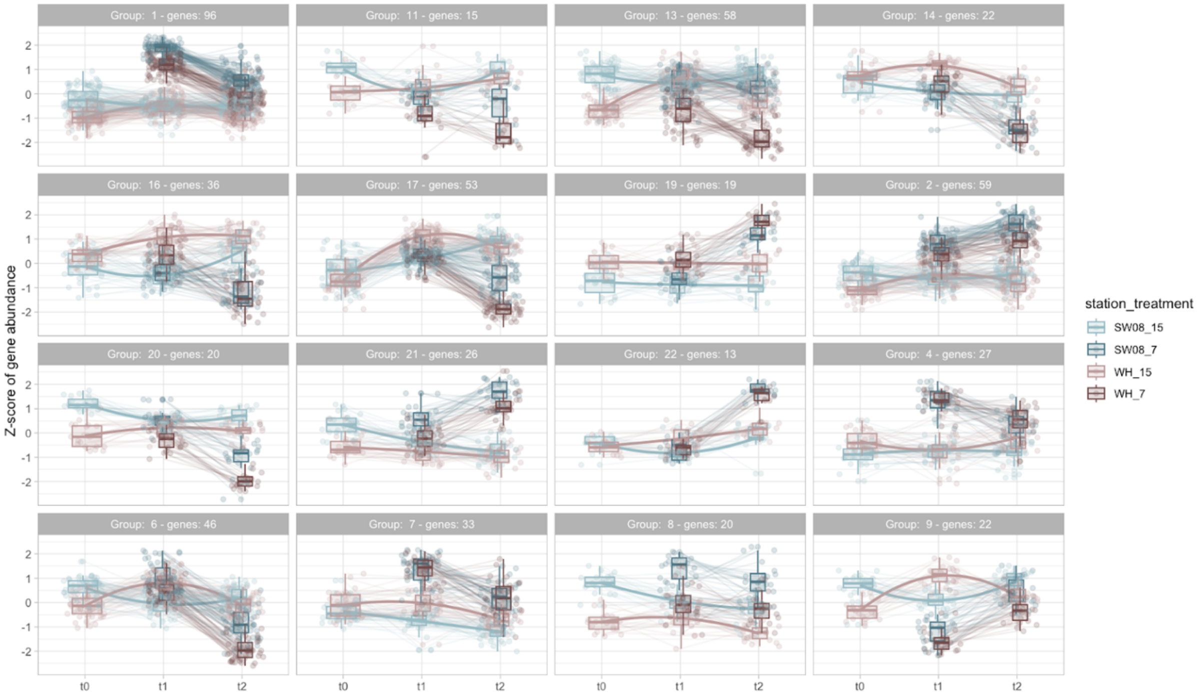


Figure S6. Differentially expressed salinity genes that show similar patterns in relative expression, clustering done using *degPatterns* from *DEGreport* (Pantano, 2022), Baltic Sea shown in blue, North Sea in red.

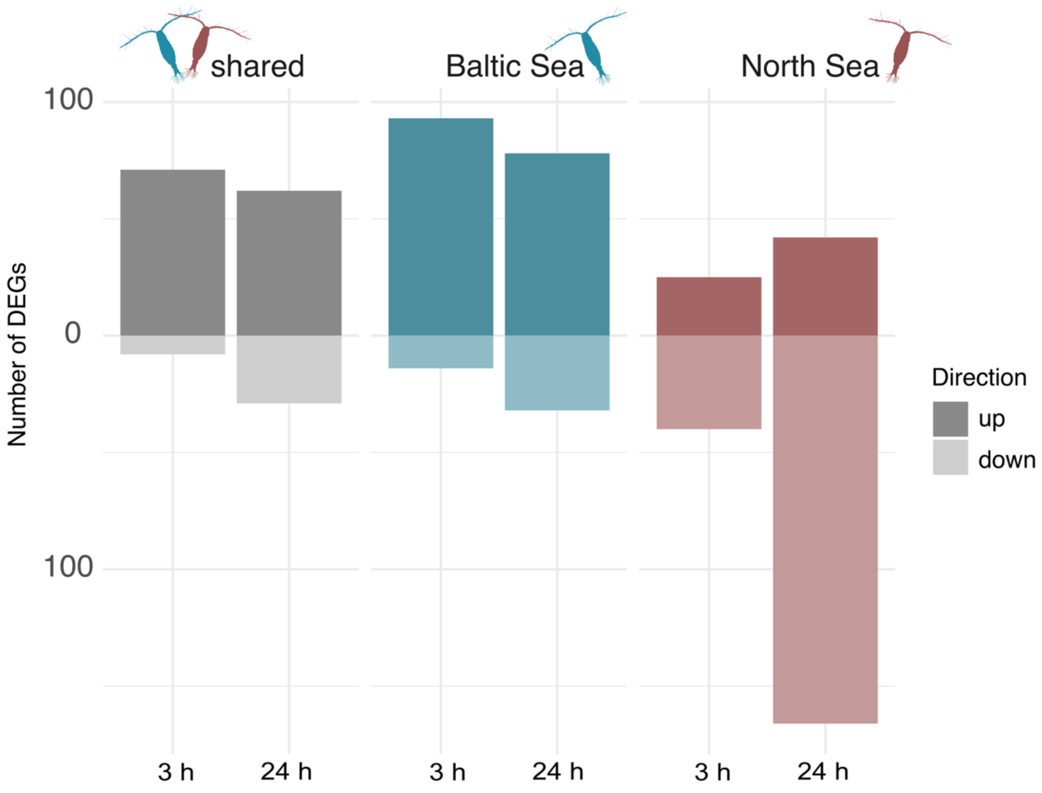


Figure S7. Differentially expressed salinity genes identified by pairwise comparisons in DESeq2 (Love et al., 2014). Over-expressed genes have positive values assigned ($\log_2$ fold change > 0, dark colors), and under-expressed genes have negative values assigned ($\log_2$ fold changes < 0, light colors). Population-specific DEGs for the Baltic Sea are shown in blue, for the North Sea in red and DEGs shared between both populations in grey.
